# Distribution and Triazole Susceptibility of Aspergilli from Clinical, Soils and Veterinary Samples in Ogun State, Nigeria

**DOI:** 10.1101/2023.12.19.572319

**Authors:** O.M. Olugbenga, T. Easter, O.B. Shittu, T.M. Obuotor, S.O. Sam-Wobo, O. Ejilude, N. van Rhijn

**Author notes:** (O.M. Olugbenga), (O.B. Shittu), (T.M. Obuotor), (S.O. Sam-Wobo), (O. Ejilude).

## Abstract

Resistance of Aspergilli to azole compounds have been on the rise globally with the majority of data coming from Europe and the Americas. However, little data from Africa is available comparing antifungal susceptibility of isolates from the environment and the clinic directly. Differences between fungal species present in the environment and their susceptibility will have a direct impact on human health and policies regarding fungal infections. Hence a One Health approach of the susceptibility of Aspergilli isolated from human, environment and veterinary samples from South western of Nigeria was carried out. Sputum, soil and animal litters/feeds samples were collected and processed using Potato Dextrose Agar (PDA) and Malt Extract Agar (MEA) supplemented with chloramphenicol to isolate *Aspergillus* species. The majority of isolates recovered were *A. niger* and *A. flavus*, with little *A. fumigatus* recovered. Susceptibility testing to voriconazole was carried out using the microdilution method results interpreted according to European Committee on Antimicrobial Susceptibility Testing (EUCAST) breakpoints. All isolates were found to be susceptible to itraconazole and voriconazole. These results show differences between species present in the environment and from patient samples compared to Europe and the Americas, highlighting a need for more fungal research focused on Africa.

## Introduction

*Aspergillus* is a genus of filamentous fungi that are ubiquitous in the environment; commonly found in soil, decaying vegetation and indoor environments. They are widely distributed and abundant in the environment and are one of the main fungal pathogens responsible for serious fungal diseases in human, animal, and degradation of plant and plant products [1]. *Aspergilli* can cause a range of infections known as aspergillosis [2, 3] which can range from mild allergic reactions to severe invasive diseases depending on the host’s underlying immunological condition. *Aspergillus* species are known to cause overlapping chronic noninvasive infection types that can range from the formation of a fungus ball (aspergilloma) to a chronic inflammatory and fibrotic process that is currently known as chronic pulmonary aspergillosis [4, 5]. It can also cause allergic bronchopulmonary aspergillosis (ABPA) and invasive pulmonary aspergillosis in immunocompromised patients [6, 7].

People living with human immunodeficiency virus/acquired immunodeficiency syndrome (HIV/AIDS), tuberculosis (TB) and other conditions that weaken the immune system have a higher risk of developing the invasive forms of aspergillosis [8, 9]. In low socioeconomic settings with inadequate diagnostic and therapeutic capabilities, pulmonary aspergillosis frequently co-infects with tuberculosis [8] and Nigeria ranks high among 30 nations with the burden for TB which is a common risk factor for chronic pulmonary aspergillosis [10]. The disease can be challenging to diagnose and requires several months of therapy. Active, drug-susceptible TB is treated with a conventional 6-month course of 4 antimicrobial medicines, and patients are supported, supervised, and given information by a medical professional or trained volunteer [11, 12]. Invasive fungal infections have been estimated to affect 11.8% of Nigerians each year [13]. 1.7 million people are thought to be affected by serious and life-threatening fungal diseases, including pulmonary aspergillosis, according to statistics from 15 of the 57 African countries [14, 15]. Given that the symptoms of aspergillosis can be mistaken for other common respiratory illnesses, health care personnel may misdiagnose patients due to a lack of awareness and education [16]. It is also possible for cases to be underreported due to a lack of systematic diagnosis and case monitoring, which results in a lack of epidemiological data and makes it challenging to determine the disease’s prevalence and devote resources for an appropriate diagnosis.

Members of the genus *Aspergillus* can also cause infections in food animals such as poultry, swine, cattle, aquaculture, and fish farming. *Aspergillus* infection (aspergillosis) and mycotoxins contamination of foods and feeds cause reduced agricultural yield, food shortage and significant economic losses as fungal growth is enhanced by the existing warm, humid atmosphere and organic substrate-rich soil in Nigeria [17, 18], this can make agricultural farms sources of *Aspergillus* infection for crops, animals, and humans alike. Different studies from Nigeria have reported *Aspergillus* sp. from cases of pulmonary TB and HIV [19-23] and soils [24-26] but reports on *Aspergilli* diversity from a One-Health approach in this low-resource setting are sparse.

Resistance to antifungal drugs is also on the increase worldwide [27] but with little information available from Nigeria, a country where serious fungal infections exist but are underreported and not well characterized. Presently, Nigeria only has four approved antifungal medications for fungal infections (amphotericin B, fluconazole, itraconazole, and voriconazole). These medications are also not easily available [28]. The agriculture sector accounts for 29.25% of Nigeria’s GDP, yet no local statistics are available about pesticide use in this industry [29]. The National Agency for Food and Drug Administration and Control in Nigeria, on the other hand, has approved the azole fungicides tricyclazole and hexaconazole. In view of the public health menace constituted by antifungal resistant *Aspergilli*, global epidemiological data and surveillance of these fungal infections are needed especially from developing African countries.

## Materials And Methods

### Study Area

The study was conducted in Ogun State which has its capital in Abeokuta. Ogun State (Fig. 1) is located in the Southwestern part of Nigeria. It lies within latitude 6°N and longitude 21/2°E and 5°E having borders with Lagos State to the South, Oyo and Osun states to the North, Ondo State to the east and the republic of Benin to the west. Abeokuta, the state capital and the largest city in the state is located at a distance of 100 km from the city of Lagos. The city is situated on the central railway line from Lagos founded in 1899, about 78 km south of Lagos. It is sited on the previous trunk road from Lagos to Ibadan.

**Figure 1.**
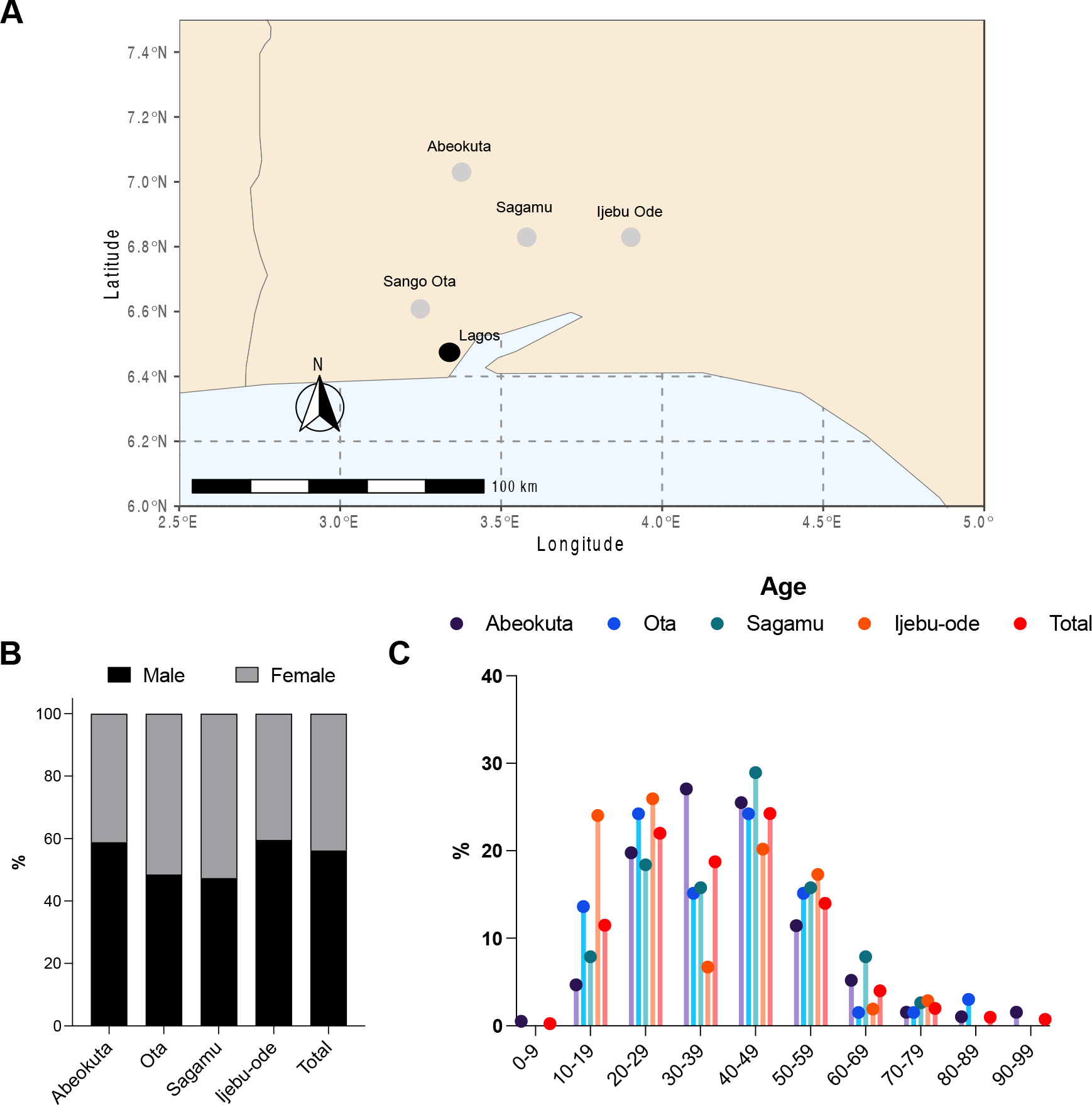
Study design and subjects. A. Study sites investigated across southern Nigeria. Sites are in grey and capital in black. B. Gender breakdown per study site. C. Age of patients assessed per study site.

### Study design

This is cross-sectional study in which sputum samples were collected from 400 TB patients between October 2020 to April 2022.

### Ethical Clearance

Ethical clearance was obtained from the Ethical Committee of the State Ministry of Health for the collection of samples from the different health facilities.

### Collection of Samples

Sputum samples from TB patients were collected from November 2020 to May 2022 from 4 state and 1 semi-private health facilities across the states (Abeokuta, Sango-Ota, Ijebu-ode and Sagamu). Samples were collected from these choice health facilities based on the high burden of TB infections. Samples from the environment (soils, animal feeds and litters) were also collected from June to July 2022. Early morning sputum was collected into sterile universal bottles from all the patients included in this study. Surface Soil samples were collected aseptically from places of much traffic into sterile universal bottles. Animal feed samples were collected from November to December 2022 randomly from bags of feed present at the time of visitation. 5 to 10g of feed were collected into sterile polyethylene bags and transported immediately to the laboratory for processing.

### Isolation of Aspergilli

Aspergilli were isolated by culturing on Sabouraud’s Dextrose Agar (HiMedia) and Malt Extract Agar (HiMedia). The culture media were prepared according to the manufacturer’s instructions and supplemented with chloramphenicol 0.05 mg/ml. All plates were incubated at 37°C for 3-5 days and examined periodically for growth development. Fungal isolates were purified according to the methods recommended [30]. The culture plates were incubated at 37°C for 96 hours after which discrete colonies were subcultured onto freshly prepared SDA slant. Isolates were characterized to a species level using distinct morphological features under a light microscope by taking a stick-tape preparation onto a drop of lactophenol blue to visualize fungal elements.

### Antifungal Susceptibility Testing

*S*usceptibility testing of the isolates to itraconazole (ITC) and voriconazole (VRZ) was performed by broth microdilution method, and results interpreted according to European Committee on Antimicrobial Susceptibility Testing (EUCAST) breakpoints and epidemiological cut-off values for species where breakpoints have not been determined [31, 32].

A working solution of the antifungal agents were prepared at 160 mg/L by diluting in sterile water and working dilutions were made starting with 160 mg/L and 1:1 serial dilution in 10% DMSO to give 80,40,20,10,5,2.5,1.25, 0.625, 0.3125 mg/L. 10 µl of the appropriate drug solution were added per well with decreasing concentration from Well 1 (High) to 10 (Low). 10ul of 10% DMSO was added to well 11. Microtitre plates were then incubated at 37 ºC for 48 hours. Plates were observed visually under the inverted microscope. The MIC is the concentration where **NO** fungal growth is observed. Alternatively, plates can be read using a plate reader OD_600nm_. For *A. fumigatus*, 48 hours is the typical endpoint.

## Acknowledgements

N.v.R and T.E. are supported by the Wellcome Trust (grant number: 226408/Z/22/Z). Other authors were self-funded. We would like to acknowledge the Ethical committee of the State Ministry of Health and all the TB local supervisors.

## Results

*Aspergillus* species are a group of filamentous fungi which are commonly found throughout the world; in various environments, soils and decaying organic matter. In this study we investigated the distribution of these *Aspergillus* species in the sputum of TB patients, soil samples and veterinary samples (animal feeds and litters) across four different zones of Ogun state, Nigeria. The study was conducted across four (4) state health facilities and a semi-private hospital in the state (Figure 1A). These sampling locations were selected because of the burden of TB patients attending the clinics. We decided to focus on patients with TB as aspergillosis and TB are commonly found together, in addition to being clinically similar,. In Nigeria, chronic pulmonary aspergillosis (CPA) has been linked to smear-negative TB and/or treatment failure [4, 23]. Nigeria as a developing country ranks first in Africa and sixth in the world accounting for about 4.6% of the global burden. The country also has a high triple burden of TB, Multi drug resistant-tuberculosis (MDR-TB) and HIV-associated TB [10].

400 sputum samples were collected from TB patients, with 11.5% having co-infection with HIV/AIDS and 10.5% were MDR-TB with a percentage of 56% being male and 44% being female (Figure 1B). Male cases of tuberculosis outnumber female cases in almost every country, which has been linked to socioeconomic and cultural impediments to healthcare access and potentially to biological differences. TB cases were the most prevalent in the age group of patients who were in the 40–49 age range (24%), followed by the 20–29 age and 30-39 age groups (22% and 18%) respectively (Figure 1C). Across the four different clinics the 10-19 age group was considerably larger in Ijebu-ode compared to other clinics, while the 30-39 age group was smaller. Sociodemographic factors seen at this clinic represent age distributions seen previously during a TB study in Ijebu-Ode. The 30-39 age group was significantly larger in the Abeokuta clinic, in line with previous reports of TB patients in Abeokuta [33, 34].

Four hundred and eighty (480) fungal organisms were recovered from the sputum of TB patients, soils and veterinary samples (feeds and litters) with 58% from sputum, 23% from the soils and 19% from the veterinary samples. The organisms recovered include *Aspergilli* (35.4%), *Mucor* (11.6%), *Rhizopus* (9.4%), *Fusarium* (0.2%), *Cladosporium* (0.2%), *Penicillium* (11%) and uncharacterized yeasts (32%). Four different *Aspergilli* were isolated; *Aspergillus niger* (59%), *Aspergillus terreus* (2%), *Aspergillus flavus* (37%) and *Aspergillus fumigatus* (1%) (Figure 2A). The result from this study is in agreement with a previous environmental survey performed in Lagos, Nigeria which showed *Aspergillus niger* and *Aspergillus flavus* as the predominant Aspergilli isolated [24].

**Figure 2.**
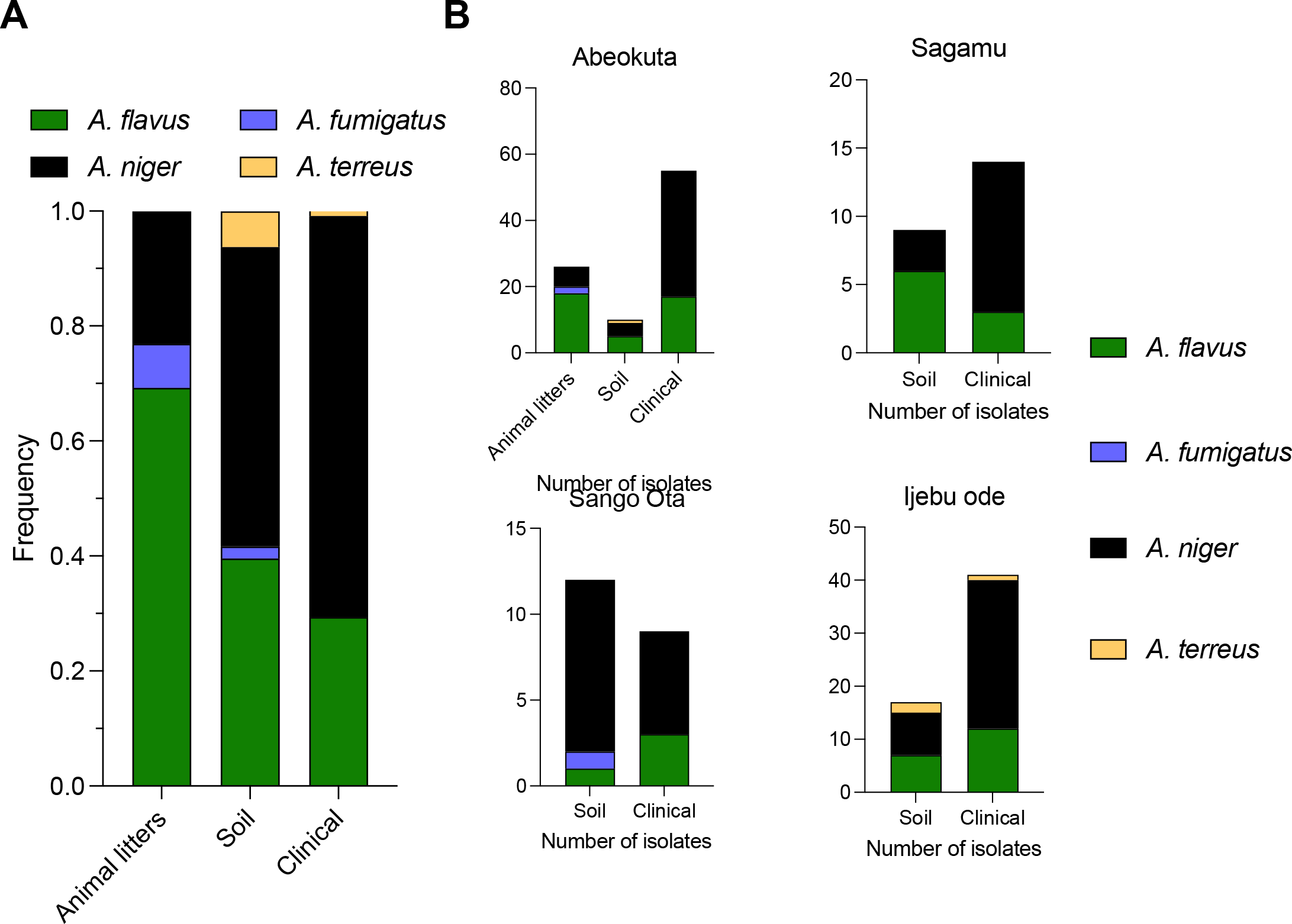
Aspergilli recovered from TB patients. A. Frequency of *Aspergillus* species obtained from samples of different sources. B. *Aspergillus* species obtained from different sites and different sample sources associated.

**Figure 3.**
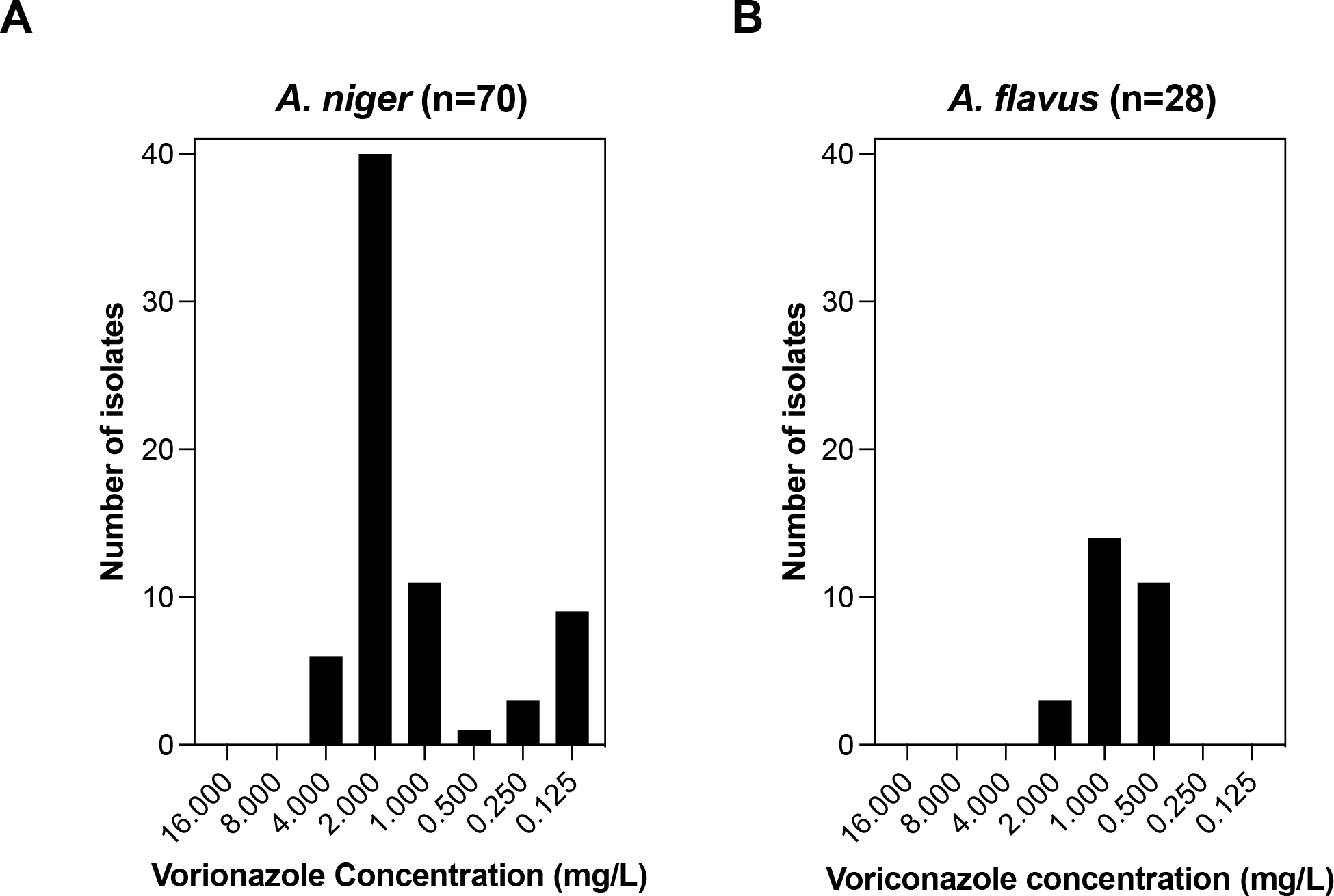
Susceptibility of Aspergillus to voriconazole. A. MIC distribution of isolates *A. niger*, MIC breakpoints were obtained via EUCAST methodology. B. MIC distribution of isolates *A. flavus*, MIC breakpoints were obtained via EUCAST methodology.

Across the different sites *A. niger* was the most predominant *Aspergillus* species isolated from clinical sputum samples (69.7%), followed by *A. flavus* (29.4%) (Figure 2A). Surprisingly, *A. fumigatus* accounted for only one isolate from a clinical sample. *A. terreus* species were isolated twice from sputum samples. This was also reflected in the soil samples were *A. niger* was the predominant *Aspergilli* isolate (52.1%), followed by *A. flavus* (39.6%) and *A. terreus* (6.25%). *A. fumigatus* was only isolated from 2.1% of soil samples. In animal litters *A. flavus* was the most isolated species (69.2%), followed by *A. niger* (23.1%) and *A. fumigatus* (7.7%).

While in animal liters from Abeokuta mostly *A. flavus* was isolated, this did not reflect the isolates from clinical samples (Figure 2B). The only site that showed more *A. niger* in the soil samples was Sango Ato, although the sample size was relatively small which could potentially account for this difference. In the Ijebu ode site *A. terreus* was identified from the soil samples in low frequency, and could also be identified from the clinical samples. Previous environmental sampling for *Aspergillus* species are in line with this finding that *A. terreus* can be found in environments across Lagos state, Nigeria [13]. In our study Ijebu-ode was the only site from which *A. terreus* could be identified.

The first line treatment for aspergillosis are the azoles [35]. Azole resistance is a significant cause of morbidity and mortality in aspergillosis patients, but data from Africa is lacking. As we isolated mainly *A. flavus* and *A. niger* from clinical samples, soils and animal litters, we sought to characterize azole resistance in Aspergilli from Ogun state, Nigeria. In this present study, we performed susceptibility testing to voriconazole for all *A. flavus* and *A. niger* isolates according to EUCAST methodology [31, 32]. For these species no official EUCAST breakpoints have been defined yet, but epidemiological cut-offs (ECOFFs) are used. The ECOFF for *A. niger* to voriconazole is 2 mg/L and for *A. flavus* 2 mg/L. Of all *A. niger* isolated tested (n=70), we found the majority to fall within the susceptible range (0.125-2 mg/L). However, 5 isolates showed a MIC of 4 mg/L, which is above the defined ECOFF. However, a two-fold difference in MIC could potentially fall within the range of technical uncertainty, so we would be hesitant to call these isolates true resistant. The *A. flavus* isolates (n=28) susceptibility to voriconazole ranged from 0.5-2 mg/L, with no resistant isolates found. This is all in line with previous findings from fungal sampling efforts in Nigeria [24].

Though there seems to be no resistance of the clinical and environmental isolates to the triazoles, there is need for continuous surveillance in Africa especially Nigeria where routine antifungal susceptibility testing is lacking. In Africa, mycology is a fascinating field full of a wide variety of fungus species that are important to human activities and the ecosystem. The distinct climatic and ecological circumstances of the continent support a diverse range of fungi, including several *Aspergilli* species. Policies that are specific to each African nation are required as a result of the distinct *Aspergillus* species that have been isolated in various areas. This is because there are significant differences in environmental conditions throughout African nations and variation within the at-risk population.

## Discussion

Fungal infections are becoming increasingly common in developing countries worldwide, partially due to more immunocompromised patients as medicine advances [1]. These patients are vulnerable to a range of fungi that can induce potentially lethal systemic infections. A variety of *Aspergillus* species are common in the environment and can cause aspergillosis in humans. TB patients are predisposed to developing aspergillosis, and is of relevance in Nigeria as TB rates are particularly high [10, 23]. This study investigated the distribution of *Aspergillus* species in sputum of TB patients, soils and veterinary samples in Ogun State, Nigeria. The age distribution revealed the highest prevalence of TB in the 40-49 age group (24.25%), which is consistent with the findings of many researchers who have linked advancing age to increased risk [36][10, 34]. Various studies also linked the majority of those with fungal infections to people in their thirtees or fourtees [21, 33]. These results are consistent throughout the centres we assessed.

*Aspergillus* species accounts for 35.7% of all the isolates from the samples. Aspergilli was the most prevalent of all the organisms isolated from all the samples with a percentage of 35.7%. The prevalence in this study is lower than that of Kumoye and Adeniji, 2023 (60.5%) but higher than that of Diba et al., 2007 (25%) and that of Mwaura et al., 2013 (2.9%) [37-39]. Four different Aspergilli were isolated; *Aspergillus niger* (59%), *Aspergillus terreus* (2%), *Aspergillus flavus* (37%) and *Aspergillus fumigatus* (1%). The result from this study showed *Aspergillus niger* to be the most prevalent of all the Aspergilli (59%) followed by *Aspergillus flavus* (37%), *Aspergillus terreus* (2%) and *Aspergillus fumigatus* (1%). While *Aspergillus niger* was prevalent in the sputum and soil samples, *Aspergillus flavus* was prevalent in the veterinary samples. The result from this study is in agreement with several environmental sampling studies from Nigeria which showed *Aspergillus niger* and *Aspergillus flavus* being predominant of all the *Aspergilli* [24, 26].

*Aspergillus fumigatus* which is one of the most common and widely studied species in the *Aspergillus* genus was however the least occurring species in all our samples. This stands in contrast to the majority of other studies in Nigeria in which *A. fumigatus* predominates, usually accounting for slightly more than half of isolates. In a study of poultry birds *A. fumigatus* accounted for 52.4% of all isolates, followed by *A. flavus* and *A. niger* [40]. Similarly, in a study in southwest Nigeria *A. fumigatus* was found in 57.1% of clinical and environmental settings [19]. However, culturing techniques vary between study (medium and temperature conditions) and the effects of sampling season and meteorological factors needs to be taken into account [41-43]. The prevalence of Aspergilli in the veterinary samples in the study was found to be 28% (25/89). This is close to the prevalence from a study conducted in Egypt which showed a prevalence of 24% [44]. Another study conducted among birds in Egypt has a prevalence of 43.2% [45]. In this study, *Aspergillus flavus* was found to be the most occurring species in the veterinary samples (20.2%). Sampling and culturing techniques vary from study to study, which further highlights the need for standardized methodologies for environmental and clinical sampling for fungi [46].

Triazole resistance, which can develop in *Aspergillus* species that are exposed to azole fungicides in the environment, is an increasing challenge in the global management of human aspergillosis [27]. Among the Aspergilli, cryptic species can exhibit intrinsic resistance to triazoles and are of particular relevance in clinical settings [47-49]. In this study, we have not assessed cryptic species via beta-tubulin sequencing and their prevalence in our samples, due to resource limitation. All isolates (both clinical and environmental) were susceptible to voriconazole, and therefore we suspect that cryptic species were not present in our samples. Our susceptibility data is in agreement with previous studies were all *Aspergilli* isolated from the soils of a south western state of Nigeria were found to be susceptible to both itraconazole and voriconazole [24]. Resistant *Aspergillus fumigatus* has however been detected in some African countries like Cameroon, Kenya and Burkina Faso [50-52]. The extensive use of triazole fungicides and their enduring presence in the ecosystem are important factors that contribute to developing and proliferation of azole resistance *Aspergillus fumigatus* [53, 54]. These environmental triazoles can decrease the number of strains that are vulnerable to azoles and promote genotypes that are resistant to them [55]. Azole containing environments may drive the emergence and expansion of azole resistant *A. fumigatus*, but are not well defined [56]. Even though from our studies all isolates were susceptible to the triazoles used, it is imperative to pay close attention to fungicide use especially in a country like Nigeria where agriculture contributes nearly thirty percent of the nation’s Gross domestic product. Antimicrobial use and resistance in farm animals as well as surveillance for resistance in environmental fungi should be monitored. Generally, systematic antifungal susceptibility studies are scarce in Africa, and more research is required to assess antifungal resistance in resource-limited settings.

